# Temperature increase affects acetate-derived methane production in Alaskan lake sediments and wetland soils

**DOI:** 10.1101/2021.08.22.457279

**Authors:** BM Dellagnezze, P. Bovio-Winkler, C. Lavergne, D.A. Menoni, F. Mosquillo, L Cabrol, M. Barret, C. Etchebehere

**Author notes:** Correspondence to: Claudia Etchebehere Microbial Ecology Laboratory, Department of Microbial Biochemistry and Genomics, Biological Research Institute “Clemente Estable”, Av. Italia 3318. CP 11600. Montevideo –Uruguay.

## Abstract

Under climate change framework, methanogens activity is expected to be strongly affected, eventually resulting in positive feedback on global climate, with higher greenhouse gas (GHG) emissions in the Arctic. This work aimed to evaluate the effect of increasing temperature on methane production rate and archaeal community of lake sediments and wetland soils from Denali to Toolik regions in Alaska (USA). For that, anaerobic acetate-amended microcosms were incubated at 5, 10, 15 and 20 °C. The acetate-derived methanogenic rate was determined and the methanogenic communities were analyzed by qPCR and 16S rRNA sequencing. Warmer temperatures yielded 4-6 times higher methane production rates and organic matter content (OM) showed significant positive correlation to methane production. Different patterns were observed in the archaeal communities after incubation at higher temperatures, with an increase in *Methanosarcina* abundance for most of the samples and *Methanosaeta* in one of the lakes tested, showing the adaptation of key acetoclastic groups among different temperatures. Our results demonstrate the impact of increasing temperature on methane production, bringing insights on key drivers involved in the process of acetoclastic methanogenic potential occurring in these ecosystems in Alaska.

## 1. INTRODUCTION

Due to polar amplification, the Arctic has been regarded as one of the most susceptible regions to the effects of climate change. The Intergovernmental Panel for Climate Change (IPCC) report (Technical Summary 2019), points out changes in Northern Hemisphere since 1950 as decrease of snow cover, shrinking of glaciers and temperature increase of permafrost. These changes are also mentioned in another report, stating since the 1970s, Alaska has warmed around 2.5°F when compared to about 1.5°F for the whole United States (NOAA, 2019).

The Arctic concentrates almost 30% of permafrost, storing around 50% of worldwide organic carbon (Zhang et al., 2003; Chowdhury, 2015). Wetlands and freshwater ecosystems (lakes, ponds) in high latitude represent one of the largest sources of the greenhouse gas methane (CH_4_), accounting for more than 40% of natural CH_4_ emissions (Rosentreter et al., 2021; Cui et al., 2015). Projections indicate that the Arctic is warming faster than tropical regions and may reach up to 9 °C of mean annual air temperature in the 22^th^ century (Collins et al., 2013; Wik et al., 2016). Such increase can lead to a positive feedback on CH_4_ production, due to accelerated thawing of permafrost enabling a major organic carbon degradation, causing longer ice-free conditions (Koven et al., 2001; Aronson et al, 2013; Basiliko et al., 2013; Cui et al., 2015; Wik et al., 2016, Wrona et al., 2015; Wen et al, 2017).

Methanogenic archaea can produce CH_4_ through different pathways from H_2_/CO_2_, methyl groups or acetate. Acetate is formed from decomposition of organic matter (OM) and is in the main CH_4_ precursor. The fate of acetate in a methanogenic environment is largely dependent on the physicochemical conditions, such as temperature, pH and salinity for example (Stams et al., 2019). In wetlands, thawed permafrost and low temperatures lakes sediments, acetate driven methanogenesis is considered to be the dominant pathway, while H_2_/CO_2_ pathways often occurs at acidic condition and mainly in deep layers (Coolen & Orsi, 2015, Cui et al., 2015, Lyu et al., 2018, Kotsyurbenko et al., 2019, Lavergne et al., 2021). Whatever the pathway considered, the last step of this process is mediated by a single enzyme, the methyl-coenzyme M reductase (mcr), reported to be highly sensitive to temperature. The α subunit of this enzyme (mcrA) is considered as a molecular marker to characterize methanogenic communities (Steinberg & Regan, 2009; He et al., 2015).

In Alaska, lake sediments and wetlands soils mainly from North Slope, like Utqiagvik (former Barrow and surroundings) and Arctic Foothills have been investigated regarding CH_4_ emission potential as well the microbial diversity involved. Lake sediments were reported regarding methane cycling using incubations (Bretz & Whalen, 2014), temperature effect on production and methane natural emission/ incubation (Lofton et al., 2014; Matheus Carnevali et al., 2015; de Jong et al., 2018). Microbial diversity involved in methanogenesis from wetland soils were evaluated in microcosms (Wagner et al., 2017; Chowdhury et al., 2015; Yang et al., 2018). Apart from Toolik region (edge of Toolik lake, within North Slope region), the other ecosystems investigated here are situated in central Alaska, near Fairbanks.

The four lakes investigated here have been previously studied, regarding the *in-situ* measurement of CH_4_ emission fluxes (Walter Anthony & Anthony, 2013, Greene et al., 2014; Sepulveda-Jauregui et al., 2015) and CH_4_ oxidation potential through incubations (Martinez-Cruz et al., 2015; He et al., 2015). Nonetheless, methanogenic communities involved in CH_4_ generation are scarcely reported and still unclear (Coolen & Orsi, 2015). Thus, this work presents novel insights on acetate derived methanogenic potential and archaeal communities involved in this process under increasing temperatures in five lake sediments and wetland soils from Alaska.

We aimed to answer the following questions: *i)* What are the effects of increasing temperature on acetoclastic methanogenic activity and archaeal community composition? *ii)* What are the differences of archaeal community structure among sampling sites *in situ* and after incubation? *iii)* Which factors affect the CH_4_ production rates? *iv)* Is the abundance of *mcrA* genes related with methanogenic activity?

## 2. MATERIAL AND METHODS

### 2.1 Sampling sites

Lake sediments (L) and wetland soil (W) samples were collected during summer 2016 in five different locations in Alaska State, USA: Killarney Lake (L1 and W1), Otto Lake (L2), Nutella Lake (L3), Goldstream Lake (L4), and Toolik Lake (W2). For lake sediments, sampling was performed at the center of the lake at maximum depth in duplicate at some meter of distance. Sample site characteristics are summarized in Table 1.

**Table 1.** Characteristics of studied ecosystems in Alaska, US.

| **Ecosystem** | **Samples** | **Coordinates** | **Location** | **Biome** | **Max Depth**<br>**(m)\*\*** |
| --- | --- | --- | --- | --- | --- |
| **Lake sediments** | L1 | 64.869879 N, 147.901824 W | Killarney lake | Boreal forest | 2.1 |
|  | L2 | 63.846202 N 149.036019 W | Otto lake | Tundra | 3.1 |
|  | L3 | 63.214548 N, 147.677642 W | Nutella lake * |  | 9.4 |
|  | L4 | 64.915801 N, 147.847705 W | Goldstream lake* | Boreal forest | 3.3 |
| **Wetland soils** | W1 | 64.86975 N, 147.900616 W | Killarney lake | Boreal forest | — |
|  | W2 | 68.62497 N, 149.600381W | Toolik lake | Tundra | — — |

\* Informal lake names.
\*\* Source: Sepulveda-Jauregui et al., 2015; Barret et al., 2019 (doi.org/10.15468/hhkhz2)

Sediment samples collected using an Ekman dredge were placed in sterile plastic bottles (1L, Nalgene®). Wetland soils were collected in 3 points and the sampling spots were divided in two subsamples, according to the depth: top (between 0-10 cm,) and bottom (10-20 cm). The average among replicates are presented aiming an overall characterization of each ecosystem (Table 2). Soil samples were collected in sterile plastic bags and placed in a cool container (4°C) before being processed, subsequently subdivided for physicochemical and molecular analyses. Subsamples for anaerobic incubations were transferred into a sterile plastic bag under N_2_ flux and sealed.

**Table 2.** Physicochemical parameters measured in lake sediment and wetland soil samples. OM (organic matter), DOC (dissolved organic carbon), DW (dry weight). The values are mean of sampling points, (field duplicates) for lakes (L1, L3,L4) and three points for wetland soils (W1, W2; T-top; B-bottom). ± indicate standard deviation

| SAMPLES | Depth (cm) | Temperature* (°C) | pH** | OM/FW (g) | OM/DW (g) | DOC (mg.L<sup>-1</sup>) | conductivity (mS/cm) | Nitrate + Nitrite (NO<sub>x</sub>) mg.L<sup>-1</sup> | Sulfates (SO<sub>4</sub>) mg.L<sup>-1</sup> |
| --- | --- | --- | --- | --- | --- | --- | --- | --- | --- |
| **Killarney (L1)** | 0-10 | 10.11 ± 0.17 | 7.07 ± 0.01 | 0.04 ± 0.01 | 0.09 ± 0.01 | 21.80 ± 0.19 | 1.73 ± 0.02 | 0.02 ± 0.001 | 2.18 ± 0.52 |
| **Otto (L2)\*\*\*** | 0-10 | 13.92 | 6.92 | 0.03 | 0.140 | 3.00 | 1.130 | 0.02 | 12.36 |
| **Nutella (L3)** | 0-10 | 12.42 ± 0.14 | 6.14 ± 0.17 | 0.02 ± 0.01 | 0.17 ± 0.11 | 4.84 ± 2.99 | 0.47 ± 0.29 | 0.02 ± 0.009 | 0.07 ± 0.01 |
| **Goldstream (L4)** | 0-10 | 19.95 ± 0.31 | 6.8 ± 0.09 | 0.02 ± 0.01 | 0.17 ± 0.07 | 12.25 ± 0.33 | 1.59 ± 0.49 | 0.01 ± 0.01 | 4.17 ± 3.71 |
| **Killarney (W1 T)** | 0-10 | 9.04 ± 2.39 | 7.12 ± 0.08 | 0.08 ± 0.02 | 0.31 ± 0.07 | 16.92 ± 5.15 | 1.03 ± 0.92 | 1.33 ± 0.04 | 0.439 ± 0.18 |
| **Killarney (W1 B)** | 10-20 | 4.15 ± 1.20 | 7.23 ± 0.01 | 0.05 | 0.090 | 21.05 ± 0.76 | 0.32 ± 0.36 | 0.99 ± 0.14 | 0.370 ± 0.09 |
| **Toolik (W2 T)** | 0-14 | 8.7 ± 0.28 | 6.21 ± 0.08 | 0.09 ± 1.56 | 0.70 ± 0.07 | 5.65 ± 2.19 | 0.21 ± 0.04 | 0.25 ± 0.007 | ND |
| **Toolik (W2 B)** | 14-20 | 3.9 | 6.37 | 0.11 | 0.71 | 7.16 | 0.14 | 0.11 | ND |

\*measured *in situ* (at the location in the moment that the sample was taken)
\*\*measured after extraction in water (1:4 w/w)
\*\*\* Otto Lake presented only one replicate

### 2.2 Physicochemical Analyses

Physicochemical parameters such as temperature, pH or conductivity were determined for each collected sample, following protocols described in Lavergne et al., (2021). Replicates of sediment/wetland soil samples were dried overnight at 105°C to determine dry weight (DW) and then heated to 550 °C for approximately three hours in a muffle furnace in order to determine organic matter content (OM). Porewater was filtered from sediments according to Jones and Willett (2006) in order to determine anions and cations concentrations. For that, an aliquot of 40 g of sample was suspended in 200 mL deionized water using a magnetic stirrer at room temperature for 1h. Then, the liquid phase was filtered through a microRhizon sampler (Rhizosphere, Netherlands) and pH was measured (HI9829, Hanna). Ammonium, nitrite, nitrate and sulfate concentrations were determined through liquid chromatography (HPLC Dionex, USA) (Table 2).

### 2.3 Microcosms set-up

A total of four lake sediments (L1 to L4) and two wetland soils (W1 and W2, including subsampling points, top and bottom) were investigated regarding methane production. Microcosms were prepared under anaerobic conditions (Pyramid GloveBox, Captair, France) in sterile glass flasks (12mL); four replicates per sample per temperature), sealed with rubber stoppers and aluminum crimp. An aliquot of 8 g OM L^-1^ of soil or sediment sample was added to each vial and the flasks were filled until 10 mL with mineral media (modified from Shelton & Tiedje, 1984), amended with acetate (30 mM), leaving 1 mL for headspace. The pH of mineral media was adjusted to the pH value of each sample measured *in situ*. All replicates were incubated in controlled temperature chambers at 5 °C, 10 °C, 15 °C and 20 °C (± 1°C). After sealing, an equilibration period of 24 hours was applied, and then flasks were purged. The acetate consumption was checked at the end of the incubation through GC-FID (Adorno et al., 2014) (data not shown).

### 2.4 Acetoclastic methanogenic activity

Accumulated biogas was monitored by measuring the pressure in the microcosm headspace, with a manual manometer (Sper Scientific, USA, Portable Handheld Pressure Meter - Wide Range, model 840065). Measurements were performed weekly or twice a week, depending on the activity of each sample. Gas chromatography (GC) was used to measure methane concentration in the headspace. An aliquot of 250 μL of headspace was injected using 0.5 mL sample-lock syringe (Hamilton, model 1750SL, USA) into a gas chromatographer (SRI Instruments, model 310C; injector temperature 40 °C, column temperature 150 °C, detector temperature 40 °C) equipped with a thermic conductivity detector (TCD) and Packed Column (6’ X1/8” SRI Instruments, USA), using argon as carrier gas (99.9% Linde, Uruguay). Calibration curves were performed periodically in gas-tight propylene bags (FlexFoil PLUS, USA) using pure methane gas (CH_4_: 99.9% Linde) and nitrogen (N_2_: 100%, Linde gas) and methane standard was injected at each measurement series for accurate measurements. Results were exported from software Peak Simple (version 3.93). Accumulated methane in both gaseous and liquid phases was calculated using the ideal gas equation and Henry’s law. Methane production rate (MPR) was calculated by linear regression, as the maximum slope of methane accumulation kinetics, following Lavergne et al. (2021). Methane production rates were normalized by the quantity of sample (calculated as dry weight-DW) in each incubation (µmol CH_4_ g^-1^ DW d^-1^). We considered the latency period as the time to reach 10% of maximum observed methane production rate.

Q_10_ coefficient was calculated according to Sepulveda-Jauregui et al (2018) to evaluate the influence of temperature on MPR. Calculation was made between 5 °C and 15 °C (Q_10_[5-15 °C]) and 10 °C and 20 °C (Q_10_[10-20 °C]). Temperature dependence of methanogenesis was also estimated using activation energy (Ea) value, applying Arrhenius equation, as described in Lavergne et al. (2021) (Table 3).

**Table 3:**
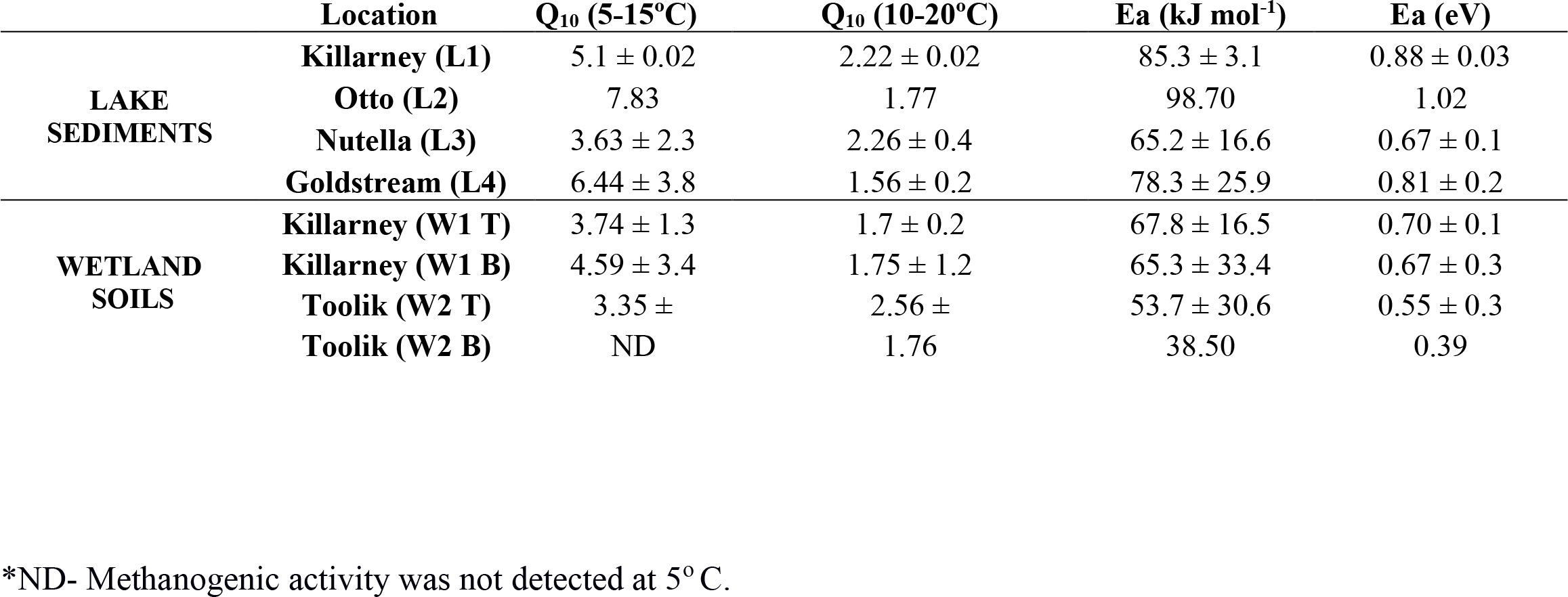
Average Q_10_ coefficient and energy activation (Ea) of methane production rate, observed for lakes sediment (L1-L4) and wetland soils (W1 and W2). Depth of wetland soils is indicated by T (top) and B (bottom).

### 2.5 DNA extraction

DNA was extracted from an aliquot of 0.25g of all *in situ* samples (lake sediments and wetland soils) and samples collected after the methanogenic assays. Samples were centrifuged (6000 rpm, 30 minutes at 4°C) at the end of the methanogenic activity (considered when no increased pressure was observed and kinetic curve reached a plateau). Power Soil Isolation kit (QIAGEN®, USA) was used following the instructor’s guide. The final DNA concentration as well as DNA purity (ratio 260/280) were quantified using Nanodrop® (Thermo Scientific, USA).

### 2.6 Abundance of targeted *mcrA* and archaeal 16S rRNA genes by qPCR

In order to quantify the archaeal 16 rRNA gene and the functional gene *mcrA* (indicative of methanogenesis), quantitative PCR (qPCR) was employed only for samples collected *in situ.* After optimization of qPCR conditions to avoid inhibition due to the sample template (data not shown), the reaction mix contained 0.4 or 1 µM of each primer (see Supp. Table 1), 2 µL of DNA template 10- or 100-fold diluted in ultrapure water (corresponding to 0.5-4 ng sample DNA), 10 µL of master mix (2X Takyon ROX SYBR master mix, Eurogentec, Belgium) and nuclease-free water to a final volume of 20 µL. All reactions were done in duplicate using AriaMX thermocycler (Agilent, USA) with Aria Software v1.3 (Agilent, USA). Standard curves and qPCR conditions are described in Supplementary material.

### 2.7 Archaeal community composition determination by 16S rRNA gene sequencing

Amplicon massive sequencing of the V3-V5 region of the 16S rRNA archaeal gene was performed using an Archaeal-specific primer set (340F 5’-CCCTAHGGGGYGCASCA-3’ and 787R 5’-GGACTACVSGGGTATCTAAT-3’) (Yu et al., 2005; Pinto et al., 2012). Only one representative sample was selected of each different incubation for sequencing. For wetland soils, only top layer samples were considered (W1T and W2T). Amplicons were checked on a 1% agarose gel electrophoresis and then purified using a commercial kit (ZR Zymoclean™ Gel DNA Recovery Kit, USA). Sequencing of the 16S rRNA gene amplicon libraries was carried out by the Ion Torrent Personal Genome Machine (PGM) platform at the Biological Research Institute “Clemente Estable” (Montevideo, Uruguay).

### 2.8 Bioinformatics analysis

Sequence analysis was performed following the ‘quantitative insights into microbial ecology’ pipeline (QIIME2 2019.10 release) (Bolyen et al., 2019). Multiplexed single-end sequencing reads (3,916,062 in total) were imported into the QIIME2.

The ‘divisive amplicon denoising algorithm’ DADA2 (Callahan et al., 2016) plugin in QIIME2 was used to filter out noise and correct errors in marginal sequences, remove chimeric sequences and singletons and dereplicate the sequences, resulting in high resolution amplicon sequence variants (ASVs). ASVs represent, as closely as possible, the original biological sequence of the sequenced amplicon.

Sequencing resulted in 790,220 sequences ranging from 976 to 72,831 per sample representing 1,289 ASVs (sequence length of 410 pb) and were classified with a classify-sklearn trained against the most recent SILVA 16S rRNA gene reference 138.1 database (Quast et al., 2012). Raw sequence data are available in the Sequence Read Archive (SRA) of NCBI under the BioProject PRJNA666194.

### 2.9 Statistical Analyses

All the statistical analysis was performed in R v3.5.1 (R Core Team, 2013) with R Studio environment (v1.3.1093). Because of non-normal distribution of the data (evaluated by Shapiro-Wilk’s test), Spearman correlation tests were performed to verify the relationship between abundance of *mcrA* genes or physicochemical parameters with MPR (methane production rate). Pearson correlation was applied to verify the effect of temperature on MPR and latency period. Significant differences and correlations were considered at p < 0.05.

In order to correlate physicochemical data with MPR, a multivariate principal component analysis (PCA) was carried out (‘FactoMineR’ package) (Husson et al., 2013). The PCA was built with 11 variables, including MPR at 5, 10, 15 and 20 °C, environmental temperature and pH (measured *in situ*), dissolved organic carbon (DOC), organic matter (OM), electric conductivity and sulfate and nitrate + nitrite concentration.

One-way ANOVA was applied to verify significant differences among MPR at four different temperatures for all samples, as well as the depth for wetland soils. The effect of temperature on MPR and latency period (lag phase) were evaluated likewise.

Regarding diversity analyses, the biom file from QIIME2 was imported and analyzed through phyloseq-modified workflow (McMurdie and Holmes, 2013). Alpha diversity in each sample was calculated based on the number of observed ASVs using Shannon diversity index after rarefying, normalized with 976 sequences per samples (Shannon, 1948; Simpson, 1949). Taxon relative abundance bar charts were generated using custom R scripts and ggplot2 (v3.3.2). Heatmap of the relative abundance of ASVs (represented at phylum and genus level) was generated using ampvis2 (v.2.6.5) (Andersen et al., 2018). Venn diagram (http://bioinformatics.psb.ugent.be/webtools/Venn/) was used to determine the shared ASVs among samples at the lowest and highest temperature in microcosms (5 °C and 20 °C). For Venn diagram construction we only considered 183 ASVs with a relative abundance above 1% of archaeal community.

Permutational analysis of variance (PERMANOVA) was applied in order to analyze the influence of ecosystem and the effect of temperature incubation on the archaeal diversity distribution, (adonis function, ‘vegan’ package; Oksanen et al., 2013). These analyses were run in two conditions: *i)* incubated + *in situ* samples; *ii)* only incubated samples at four different temperatures. Principal coordinate analysis (PCoA) was used to evaluate the distribution of samples only in natural condition (*in situ*). PCoA using Bray Curtis distance metrics of non-transformed relative abundance table was computed with phyloseq and plotted using ggplot2.

## 3. RESULTS

### 3.1 Methane production rate (MPR) in incubated lake sediments and wetland soils

Methane production was monitored along time in the incubations at the different temperatures.

Temperature influenced significantly MPR and latency periods in all lake sediments and wetland top samples. Pearson correlation showed strong correlation between temperature and MPR, as temperature and latency period (p < 0.05, ρ > 0.9). All lake sediment samples incubated at warmer temperatures (15 °C and 20 °C showed significant higher MPR (p < 0.05) and longer latency periods (p < 0.05) when compared to the coldest temperatures (5 °C) (Fig. 1A and 1B).

**Figure 1.**
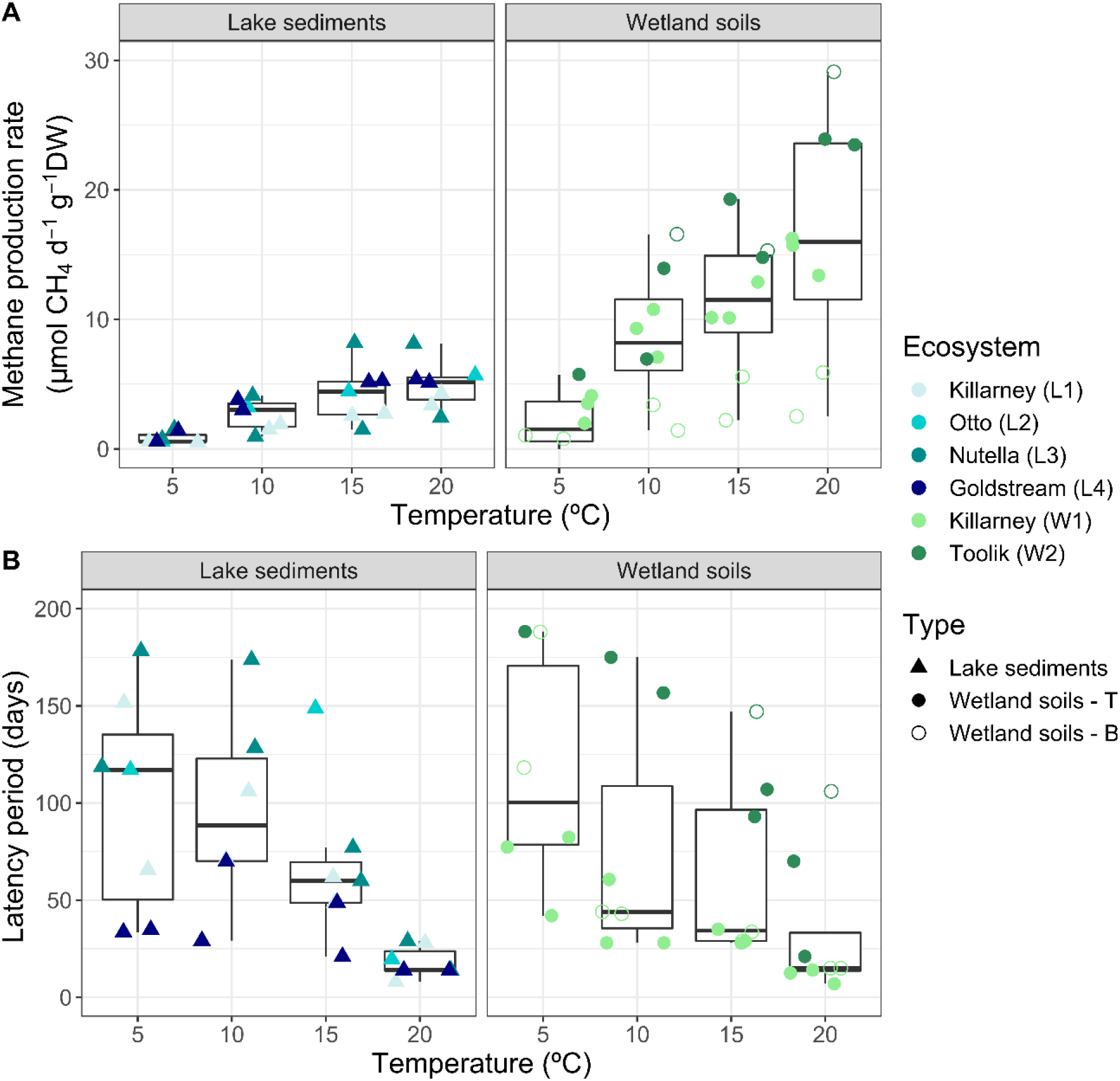
Boxplot representing the influence of temperature on both CH_4_ production rates (panel A) and latency periods (panel B) in acetate-amended microcosms from lake sediments and wetland soils. Type of freshwater systems was plotted by different shapes and the 6 ecosystems were represented by colors. For wetland soils, distinction was made between top (T) (disks) and bottom (B) (open circles) samples.

The rate of acetate-driven methane production from lake sediment samples was higher at 20°C and varied from 3.8 ± 0.03 (Killarney Lake) to 5.6 ± 0.6 µmoles CH_4_ g^-1^ DW d^-1^ (Otto Lake) (Fig. 1). At 5°C, MPR were on average 5 times lower than at 20 °C, ranging from 0.54 ± 0.05 (Killarney Lake) to 1.15 ± 0.2 µmoles CH_4_ g^-1^ DW d^-1^ (Nutella Lake). Despite significant differences of MPRs among four temperatures (5 to 20°C), when compared among the four lake sediments (L1-L4) no significant differences were found (p > 0.05) (Fig 1A).

As lake sediments, temperature had significative and strong positive correlation with MPRs (p < 0.05, ρ > 0.9) in all wetland soil samples, exceptionally to Toolik B. Pearson correlation between temperature and latency periods was statistically significant and strongly negative (p < 0.05, ρ > - 0.9) only for top samples (W1T, W2T). MPRs were higher at 15 °C and 20 °C than at 5°C (p < 0.05), which at 20 °C MPR increased about 4-5 times ranging from 4.21 ± 0.4 (Killarney - W1B) to 29.12 ± 0.7 (Toolik W2B) µmoles CH_4_ g^-1^ DW d^-1^. At 5°C, methane production was not detected for the bottom layer of the Toolik (W2 B) and the highest MPR were observed at Toolik T (5.7 ± 1.8 µmoles CH_4_ g^-1^ DW d^-1^. Comparing both ecosystems, MPRs showed no significant differences (p > 0.05) between Killarney and Toolik soils. Nonetheless, depth had significative influence on MPR only in Killarney wetland soil (W1 T ≠ W1 B) (Fig. 1A). Latency periods pronouncedly diverged in top layer samples (p < 0.05, W1T and W2T), which Killarney (W1 T) presented shorter latency period, (10 days at 20 °C to 68 at 5 °C) than samples from Toolik (W2 T) with 45 days at 20 °C to 146 at 5 °C) (Fig. 2A).

**Figure 2.**
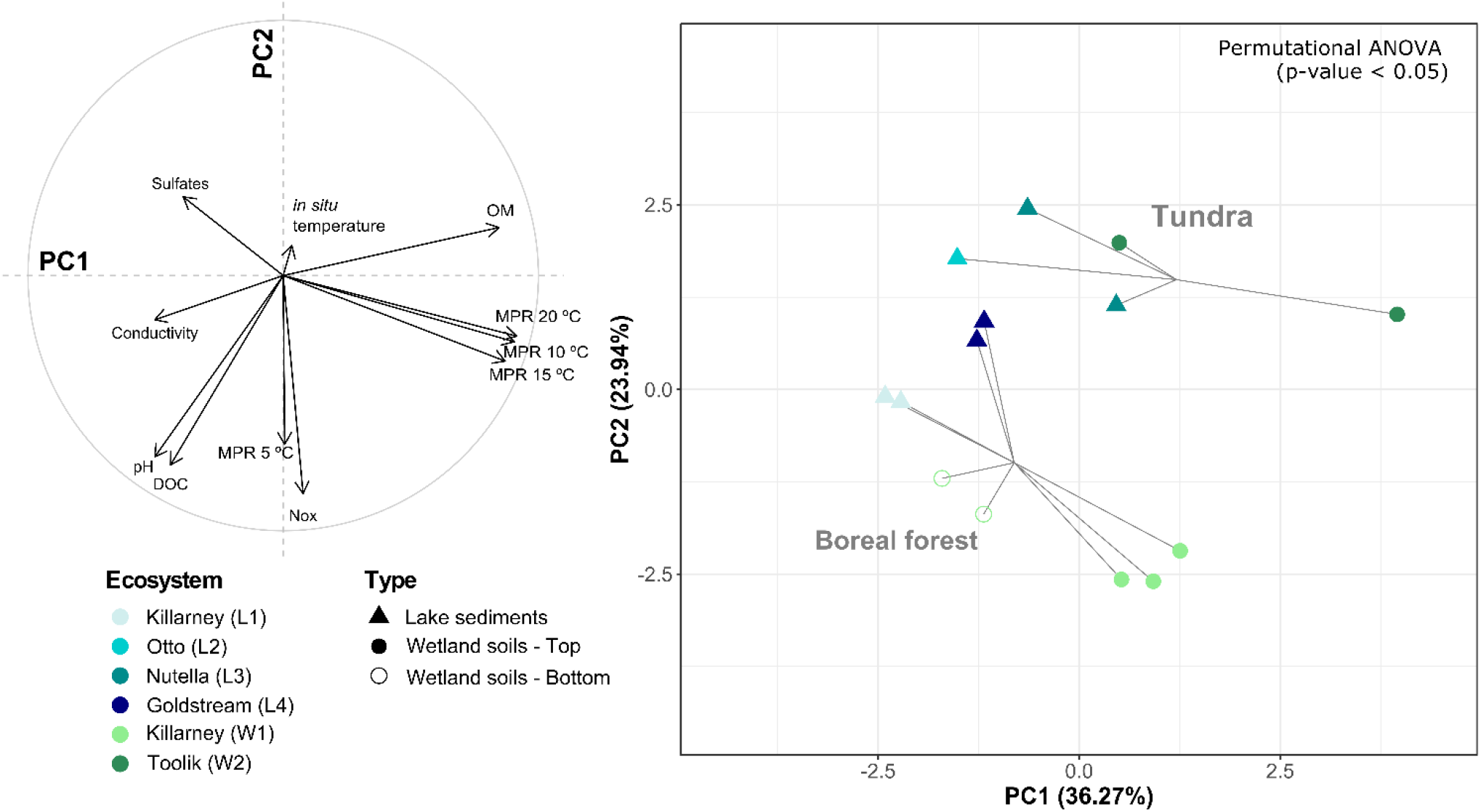
Principal component analysis (PCA) built from 11 variables showing the correlation between CH_4_ production rates (MPRs measured at 5, 10, 15 and 20 °C) and *in situ* physicochemical data. Left panel (A) showed the correlation plot of variables and right panel (B) showed the site position on the ordination colored by ecosystem, while the symbol shape represents the different ecosystem types. Significant segregation by biome (PERMANOVA, F = 4.18, p-value < 0.05) was plotted as spiders (95% confidence level). OM stands for organic matter and NOx for nitrates + nitrites.

For all samples, the Q_10_ indicated that an increase by 10 °C had a positive impact on methanogenic production (in average 5 ± 2.2 for Q_10_[5-15 °C] and 2 ± 0.5 for Q_10_[10-20 °C] (Table 3). It is worthy to note that an increase of 10°C had a major effect in the lowest temperature range (5 to 15 °C) compared to the highest range (t-test, t = 4.42, p < 0.05). Moreover, the overall temperature dependence of methanogenesis evaluated by the energy activation (Ea) showed similar values for lake sediment samples (mean: 81.9 ± 11.4 kJ mol^-1^, corresponding to 0.84 ± 0.15 eV) and wetland soil (mean: 56.3 ± 26.8 kJ mol^-1^, corresponding to 0.58 ± 0.27 eV) (Table 3). The highest values was displayed by the Otto Lake, reaching up to 98.7 kJ mol^-1^ (1.02 eV).

### 3.2 Physicochemical analysis and effects on methane production

Values of several physicochemical parameters measured in the samples are presented in Table 2. The highest dissolved organic carbon (DOC) contents were observed in Killarney lake sediment samples (21.81 mg L^-1^) and Killarney soil bottom samples (21.01 mg L^-1^). Also, Killarney top (W1 T) presented the highest concentration of nitrate + nitrite (0.133 mg L^-1^). Regarding organic matter content (OM), the highest values were observed in the samples from Goldstream and Nutella Lakes, 0.179 and 0.172 g OM g^-1^ DW, respectively and Toolik bottom (W2 B), 0.716 g OM g^-1^ DW. Otto lake sample presented the highest content of sulfate (12.36 mg L^-1^).

To elucidate which physicochemical factors was related to the MPRs, principal component analysis (PCA) was applied to 11 variables. The two first dimensions of the PCA built with MPRs at the four tested temperatures (5, 10, 15 and 20°C), in situ temperature, conductivity, pH, dissolved organic carbon (DOC), sulfate, nitrate + nitrite concentrations and OM represented 60.21% of the total dataset variation (Table 2, Fig. 2A). OM concentration was significantly correlated to MPR in all samples at 10, 15 and 20 °C (Spearman correlation, ρ > 0.6, p-value < 0.05). Moreover, based on the two first dimensions of the PCA, samples were significantly clustered according to their biome, with a clear distinction between tundra (Toolik soil and Nutella Lake) and boreal forest (Killarney, Goldstream) (PERMANOVA, F = 4.18, p-value < 0.05) (Fig. 2B).

### 3.3 16S rRNA archaeal and *mcrA* gene abundance in sediment lake and wetland soils

In all *in-situ* samples, both archaeal communities and methanogens were detected (Fig. 3). Among lake sediments, the highest archaeal 16S rRNA gene copy number was observed for samples from Goldstream lake (L4) (1.05 x 10^9^ ± 2.72 x 10^7^ copies g^-1^ DW) and the lowest value was observed in Killarney lake (L1) (4.51 x 10^7^ ± 2.51 x 10^6^ copies g^-1^ DW). For wetland soils, the highest and lowest values of archaeal 16S rRNA gene copy number were observed respectively in bottom samples from Toolik (W2 B) (2.07 x 10^9^ ± 3.96 x 10^6^ copies g^-1^ DW) and Killarney (W1 B) (6.14 x 10^7^ ± 3.17 x 10^6^ copies g^-1^ DW).

**Figure 3.**
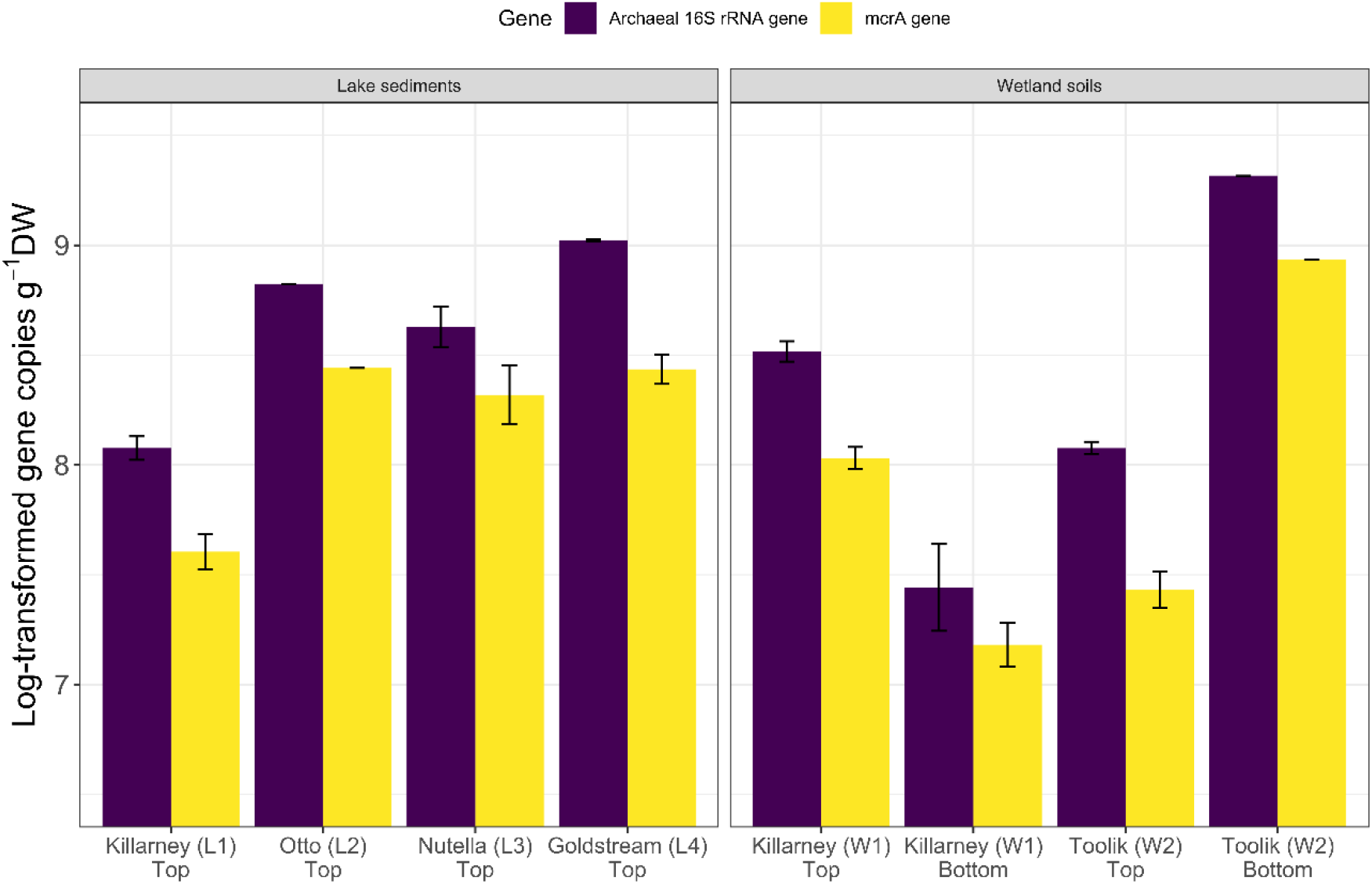
Abundance of *mcrA* gene (in purple) and archaeal 16S rRNA gene (in yellow) quantified by qPCR from *in situ* lake sediment (L1-L4) and wetland soil (W1, W2, top and bottom). Error bars represented standard error between field replicates.

**Figure 4.**
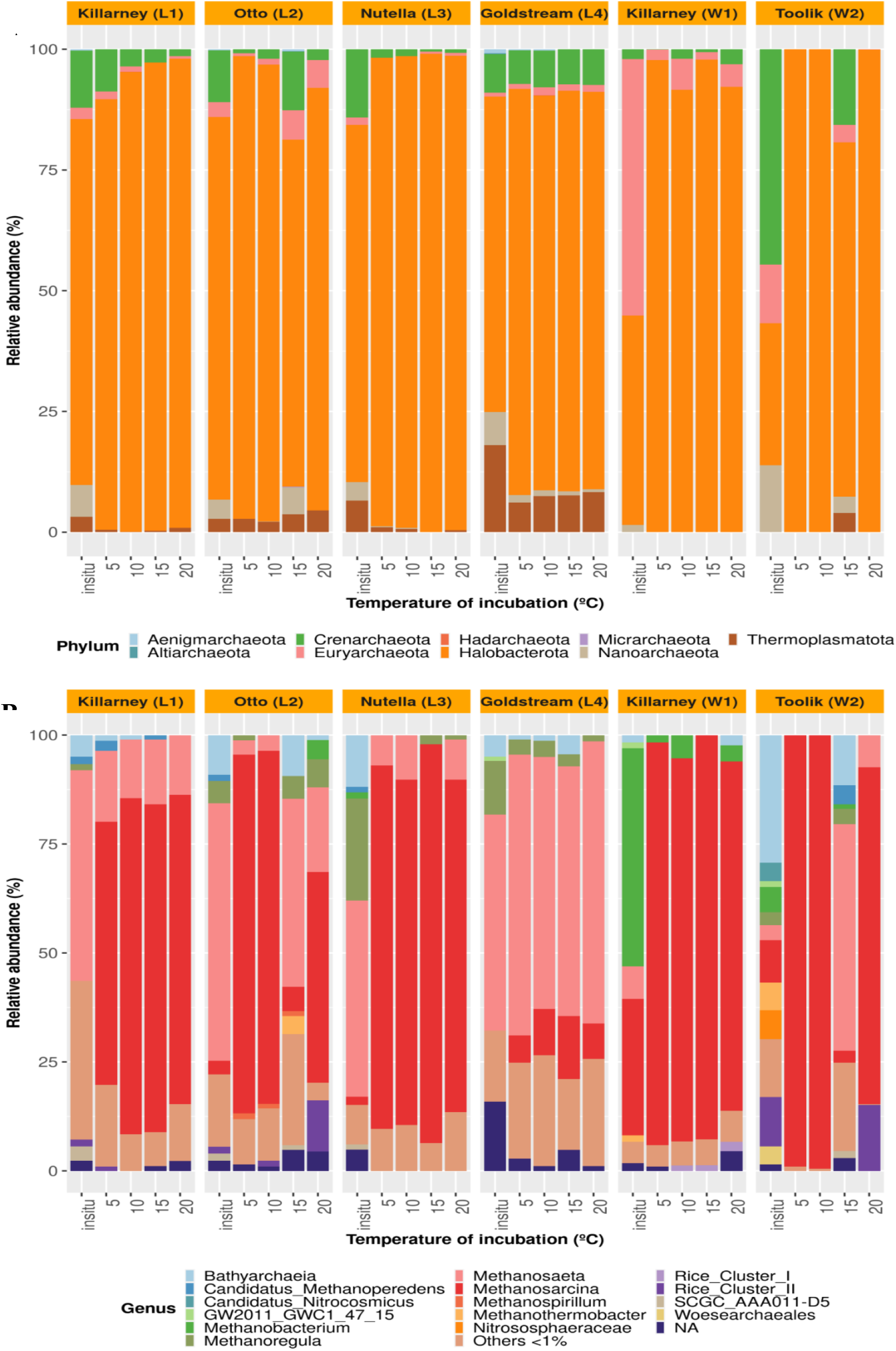
Taxonomic composition of the archaeal communities at the phylum (A) and genus (B) level, from *in situ* samples and after microcosm incubations at four temperatures, from four lake sediments and two wetland soils, according to 16S rRNA gene sequence analysis. Sequences with no taxonomic assignment (uncultured or unclassified) were grouped as NA. Other ASVs with less than 1% are indicated as “Others <1%)

The highest abundance of functional gene *mcrA* was found in Nutella Lake samples (L3) (2.81 x 10^8^ copies g^-1^ DW, with 51% of *mcrA* in total Archaea) and Goldstream Lake samples (L4) (2.96 x 10^8^ copies g^-1^ DW, 28% of *mcrA* in total Archaea). Likewise, among wetland soils, the highest *mcrA* abundances were observed in Toolik bottom soil samples (W2 B) (8.59 x 10^9^ copies g^-1^ DW, 41% of *mcrA* in total Archaea), followed by top soil samples from Killarney (W1 T) (1.15 x 10^8^ copies g^-1^ DW, 33% of *mcrA* in total Archaea). No significant correlations were found between *mcrA* gene abundances and MPRs independently of the considered temperature (Spearman correlations, ρ = −0.08 - 0.03, p-value > 0.05).

### 3.4 Methanogenic community analysis

Archaeal communities were largely dominated by the phylum Halobacterota (36.1 % of total Archaea) in all incubated samples at all temperatures, and in most *in-situ* samples, followed by phyla Crenarcheota, Euryarcheota, Thermoplasmatota and Nanoarcheota (Fig. 5A). The characterization at the genus level, revealed the presence of known methanogens, representing 26 % of the total community.

In *in situ* samples, *Methanosaeta* was the most abundant genus in all lake sediment samples (ranging from 45 to 60%, representing 6.2 % of the total Archaea) while *Methanoregula* was found above 10% only in Nutella and Goldstream Lakes samples (25 and 14%, respectively). *Methanosarcina* showed very low abundance in Killarney, Otto and Nutella Lake (1.8 to 4.5%) and was not detected in Goldstream Lake. Bathyarchaeia class ranged around 10% in all lakes. From wetland soils, archaeal diversity was assessed from the active layer (upper layer, 0-10 cm). In *in-situ* Killarney soils, *Methanobacterium* was found as the most abundant (50.8%), followed by *Methanosarcina* (31.2%). The *in-situ* community from Toolik soil (W2 T) showed the highest diversity (Shannon index, 3.66) (Supp. Fig. S4), dominated by Bathyarchaeia class (31% of the total community), followed by Rice Cluster II family (11.3%) and *Methanosarcina* (9.7%). Woesearchaeales (8.2%) and Nitrososphareaceae (8.4%) were detected more abundantly in this sample (Supp. Fig. S1).

Alpha diversity analysis (Shannon index) showed a decline after microcosm incubations in all samples, being significative lower at 5, 10 and 20 °C (p < 0.05), mainly in Toolik soil (Supp. Fig. S4). Indeed, *Methanosarcina* was globally predominant in microcosms (60-99% of archaeal relative abundance), at all temperatures, in all samples, except for the samples from Goldstream Lake, where the genus *Methanosaeta* prevailed at all temperatures (49-67% of relative abundance). *Methanosaeta* was found at 15°C incubation in Otto lake (44.7%) and Toolik peat (52.7%) (Supp. Fig. S1). The effect of different temperatures on diversity (Shannon index) was not significant in any sample (Spearman correlation, p > 0.05, ρ < 0.4). DNA extraction from samples of Otto Lake and Toolik soils for sequencing were extracted later than others, which could have infer in this variance.

The most abundant genera found in microcosms were *Methanosarcina* (represented by 97 ASVs) or *Methanosaeta* (53 ASVs). Despite being affiliated to the same genus, the most part of ASVs were observed as site-specific, exceptionally some occurring in Otto lake, punctually in determined temperature (Supp. Fig. S2). Venn diagram was constructed to evaluate the shared ASVs between samples taken *in situ* and after incubation, at the lowest (5 °C) and highest (20 °C) temperatures.

In wetland soils, only 2 ASVs (both related to *Methanosarcina*) were shared between *in situ* and incubated samples (at 5°C and 20 °C), 10 ASVs (where 5 related to *Methanosaeta*, 4 to *Methanosarcina* and one to *Methanoregula*) were shared in lake sediments (Supp. Fig. S3), showing that the incubation induced a stronger divergence from natural conditions. A total of 10 and 16 ASVs mainly related to *Methanosarcina* genus were shared between samples incubated at 5 and 20° C for wetland soils and lakes sediments, respectively, mainly affiliated to *Methanosarcina* (Supp. Table S2). This result suggests that these methanogens, key acetoclastic groups (mostly *Methanosarcina* and *Methanosaeta*) could adapt to higher temperature (20 °C), in a psychrophilic/psychrotolerant range.

PCoA and PERMANOVA based on all ASVs revealed that samples primarily clustered according to the ecosystem type (wetland soils and sediment lakes), when considering all conditions evaluated; *in situ* + incubation at different temperatures, incubated samples only (Bray Curtis distance-based PCoA, PERMANOVA, p < 0.05, (Fig. 6A & C). As a general rule, the *in situ* samples were clearly separated from the incubated ones, except for the Goldstream lake (L4) samples exhibiting a very stable archaeal composition *in situ* and after incubation (whatever the temperature) (Fig. 6A). Under natural conditions (*in situ*), lake sediments samples displayed similar archaeal community structure, with a clear cluster while wetland soils highly diverged between each other, reflecting the strongly different ecosystems between Toolik (tundra) and Killarney (boreal forest) (Fig. 6B).

## 4. DISCUSSION

### 4.1 Effects of increasing temperature and other environmental factors on CH_4_ production

Higher temperatures (15 and 20 °C) reduced latency period and strongly affected MPRs resulting in a 4-6-fold increase of acetate-based MPR when compared to the lowest ones (5 and 10 °C). It is well defined that temperature is a limiting factor, defining metabolic functions and our results corroborates with other works performed in Arctic environment using incubation experiments (Hoj et al., 2008; Lofton et al., 2015; Blake et al., 2015, Herndon et al., 2015).

The Q_10_ coefficient represents the change of a rate in 10 °C. According to Bennett (1990) and applied in Fuchs et al. (2016), Q_10_ values between 0.2 and 0.8 can be considered as negative correlation, at 0.8–1.5 not impacted by temperature and as positive correlation is stated values at >1.5. Also, Q_10_ values showed positive correlation at the lowest temperature range (5-15 °C), mainly for Otto and Goldstream Lakes (Q_10 >_ 6), showing strong temperature sensitivity, suggesting more energy input to methanogenesis than in the other ecosystems evaluated. These values can be comparable to those found in acetoclastic incubation experiments at low temperatures (de Jong et al., 2018, Blake et al., 2015). However, Q_10_ values were slightly lower than observed in Sepulveda-Jauregui et al. (2018), a study involving both lakes without substrate addition and tested in a broader temperature range. The overall methanogenesis dependence to temperature (Ea) was on average 0.84 ± 0.15 eV in lake sediments. This value fits within the universal Ea range defined by Yvon-Durocher et al. (2014) (0.82 –1.27 eV) and also corresponds to similar values recently published by Lavergne et al. (2021) using the same protocol in sub-Antarctic lake sediments.

As mentioned previously, ecosystems investigated here were studied by Sepulveda-Jauregui and collaborators (2015) through field measurements of CH_4_ emission associated with biogeographical and physicochemical features. They verified lakes Nutella and Otto as non-yedoma type peatlands, emitting lower CH_4_ amounts than Goldstream and Killarney classified as yedoma type peatlands. Despite diverging in natural conditions, in controlled anoxic microcosms all samples reached around 5 µmoles of CH_4_ g^-1^ DW d^-1^ at 20 °C. Considering the methanogenesis stimulation by 30 mM of acetate and the temperatures tested, the MPRs were lower than other reported in lake sediments from Alaska. From the North Slope region (Alaska), microcosm amended with 2 mM of acetate at 4 and 10 °C, presented a higher MPR of 4.7 µmoles CH_4_ g^-1^ DW d^-1^ at 10°C (de Jong et al., 2018). In non-amended incubation, sediment lake samples from Barrow surroundings yielded a rate of 2.2 µmoles CH_4_ g^-1^ DW d^-1^ at 10 °C (Matheus Carnevali et al., 2015), similar to our results, despite no substrate addition. Our MPRs are also almost 3-times lower than the rates found in a similar study of sub-Antarctic lake sediments stimulating CH_4_ production with 30 mM of acetate within the same temperature range (Lavergne et al., 2021).

MPRs were in average 3 times higher in wetland soil samples than lake sediments. Although not significant differences observed (p > 0.05), between two wetland ecosystems, Toolik samples yielded higher MPR than Killarney, with higher MPR in Toolik soil bottom (W2 B; 10-20 cm depth). These samples were collected from Tundra region, and presented the highest OM content among all samples evaluated, with strong correlation with MPR (ρ > 0.8). Shaver et al. (2013) reported soils ecosystems in Toolik Lake region, stating these soils typically have high OM content overlying a poorly developed mineral soil. Thus, along with temperature, substrate availability is crucial for CH_4_ cycle in natural systems. Some works performed in different locations of the Arctic report that organic rich soils can yield higher amounts of CH_4_ than mineral soils; Alaska (Chowdhury et al., 2015), Siberia (Ganzert et al., 2006), Norway (Høj et al., 2008). Depth is a factor linked to the moisture and temperature of soil, influencing the turnover of soil OM (Kechavarzi et al., 2010). CH_4_ accumulation at increased temperatures can be higher in deep layers than in surface; however, this can vary depending on the soil type (Wang et al., 2017).

### 4.2 Methanogens abundance and MPRs

The MPR values do not accurately correlate with *mcrA* gene abundances. While previous works have shown that *mcrA* gene could be a good molecular marker for methanogenesis (Lavergne et al 2021, Zhang et al., 2019, Yang et al., 2018), we found no significant correlation between *mcrA* gene abundance (*in situ* samples) and MPRs, which is in line with Freitag and Prosser (2009). Moreover, qPCR was carried out based on DNA approach, which provide us a “general” information and potential, but not the functionally active genes during incubation period. Franchi et al., (2018) evaluated *mcrA* gene in anaerobic digestion process under both approaches (DNA and RNA), observing in spite of high abundance, low gene expression was observed and no correlation with methane production was found as well.

Despite no positive correlation, the highest abundances of *mcrA* were observed in the samples taken from Goldstream Lake (L4) and Toolik soils (“bottom”-W2 B), also presenting the highest MPR. However, the detection of *mcrA* genes and the MPR obtained even at 5° C could have indicated the potential of methanogenesis in the ecosystems sampled.

### 4.3 Methanogenic microbial communities’ composition

The community composition was more influenced by ecosystem type than by temperature of incubation. When PCoA was performed using all the 16S rRNA genes sequences datasets, a separation of the samples according to the ecosystem and not according to the temperature of incubation was observed. The same separation was observed when the PCoA was performed using only the samples taken after incubation. These results indicate that methanogenic communities were particularly influenced by ecosystem type. In addition, similar patterns where other factors than temperature influence diversity were observed in wetland samples from California, USA (He et al., 2015), from lake sediment in Germany (Glissman et al., 2002), in Chinese Zoige wetland (Tibetan plateau) (Cui at al., 2015).

The communities from the samples taken *in situ* presented different composition according to the ecosystem type. The communities from the lakes were dominated by *Methanosaeta* genus *(Methanosarcinales)* (from 45.8 to 60.9% relative abundance), followed by *Methanoregula* and *Methanobacterium* genera. The samples taken from wetlands exhibited different methanogenic communities, with a dominance of *Methanosarcina* and *Methanobacterium* in Killarney wetland soil (W1) and a dominance of Bathyarchaeia and Rice cluster II in Toolik soil (W2). Our findings corroborate with other works performed in Arctic environment. *Methanosarcinales, Methanomicrobiales, Methanobacterales* orders have been reported in high abundance in response to the permafrost thaw in Tibetan Plateau and in Chinese wetland (Wei et al., 2018; Cui et al., 2015), palsa peatland in Norway (Liebner et al., 2015), and permafrost thaw ponds in Canada (Crevecoeur et al., 2016). In Alaska, *Methanosarcinales, Methanomicrobiales* were found as active members in upper layer of sediment lakes from near Barrow, North Slope (Matheus Carnevali et al., 2018) and in thawed active layer from Hess Creek (Mackelprang et al., 2011). At genus level, *Methanoregula* was found as the most abundant in sediment lake samples and *Methanosarcina* and *Methanobacterium* accounted for more than 90% of all classified archaeal methanogens in wetland ecosystems in the Canadian Arctic (Stoeva et al. 2013; Frank-Fahle et al., 2014). In Alaska, *Methanobacterium* and *Methanosarcina* significantly correlated with the methane production at various depth levels in tundra and boreal forest soils cores (Tripathi et al. 2017).

The groups of archaea mostly present in Toolik samples was also previously observed in samples from cold environments. Rice cluster II lineage have been described as dominant phylotype in thawing permafrost in Sweden (Mondav et al., 2014) and boreal peatlands in Finland (Juottonen et al., 2015). Toolik soil harbored most Bathyarchaeia members (Crenarcheota/Bathyarcheota phylum) and almost exclusively ASVs related to the recently classified Woesearchaeales order and Nitrososphaeraceae. Bathyarchaeia is a wide group, associated with anoxic environments, like termite gut (Loh et al., 2020), deep seafloor (Chen et al, 2020) and freshwater (Fillol et al., 2015). Their physiology is associated with several functions as acetogenesis and methane metabolism (Zhoug et al., 2018). Woesearchaeales (phylum Nanoarcheota, superphylum DPANN) are also reported with anaerobic metabolism in deep marine sediments, for example (Carrier et al., 2020). Woesearchaeales and Bathyarchaeia were described in thermokarst lakes in Alaska (Matheus Carnevali et al., 2018), in lake sediments from south Chile (Lavergne et al., 2021) and tundra permafrost soil in Alaska (Tripathi et al., 2019). Nitrososphaeraceae family members were related to ammonia oxidation, have been detected in tundra soil in Russia (Hetz & Horn, 2021) and isolated from peatland in Norway (Candidatus *Nitrosocosmicus arcticus*) (Alves et al., 2018). Then, most of the groups detected in our work were previously found in cold environments suggesting their adaptation to low temperatures.

After microcosms incubation, the diversity decreased and two different patterns were observed: in most samples, a shift was observed with an increase in abundance of *Methanosarcina*, while in the samples taken from Goldstream lake, the in-situ dominant *Methanosaeta* persisted and was even enriched after incubation.

In the samples from lakes Killarney (L1), Otto (L2) and Nutella (L3), similar patterns were observed each other, *Methanosaeta* prevailed in the samples taken *in situ* and *Methanosarcina* predominated after incubation. In natural environment, low concentration of acetate and low temperatures may favor the obligate-acetoclastic *Methanosaeta*, which is more competitive than *Methanosarcina* under these conditions (Stam et al.,2019; Kotsyurbenko et al. 2019). Thus, despite the previous high abundance of *Methanosaeta in situ*, the generalist feature of *Methanosarcina* to use a broad range of substrates, producing methane through distinct metabolic pathways, may explain their dominance. Moreover, compared to *Methanosaeta*, *Methanosarcina* can better tolerate high acetate concentration (Stam, et al., 2019), which could have influenced their predominance in these in acetate amended microcosm.

The samples from Goldstream Lake presented a different behavior, with high abundance of *Methanosaeta* before and after incubation. A reasonable explanation for *Methanosaeta* prevalence exclusively in Goldstream Lake might be associated with the very low abundance of *Methanosarcina* (not detected) in natural samples (*in situ*). Also, *Methanosaeta* might be metabolically active in environmental condition; once active *in situ*, associated to very low abundance of *Methanosarcina*, may have led it to surpass and thrive on other members.

Acetoclastic pathway was reported to be the most likely process in thawing permafrost and one of the most important methanogenic substrates in low temperature environments (Frank-Fahle et al., 2014, Mackelprang et al., 2011, Coolen & Orsi, 2015). Our results corroborate with other works performed through acetate incubation. *Methanosaetaceae* family was found as the most abundant in sediment lakes from Alaska with low concentration (2 mM), at 10 °C (de Jong et al. 2018). From lake sediments in Norway, Blake et al. (2015) observed this shift where *Methanosaetaceae* was the most abundant before incubation, decreasing at 5°C and *Methanosarcinaceae* and *Methanosaetaceae* compounded major part of the community at 30°C. On the other hand, our results contrast with a study in Canadian High Arctic permafrost where *Methanococcales* and *Methanomicrobiales* were found in acetoclastic incubations (30 mM as in our study) at 4 °C and 22 °C (Supplementary Table S2).

The samples from the two wetland sites presented higher diversity and in particular, the sample taken from Toolik was very diverse presenting high abundance of “not classified” sequences, or classified as uncultured microorganisms, suggesting this environment can harbor many unknown archaeal microorganisms so far. After incubation at four different temperatures, the pattern was similar to most of the other samples, with an enrichment of *Methanosarcina*.

One of the limitations of this work was that only acetoclastic methanogenic capacity was studied, then, we cannot infer from these results the response of the whole methanogens to temperature increase. In sub-Antarctic lake sediments, Lavergne et al. (2021) stimulated methanogenesis separately by acetate (30 mM) and H_2_/CO_2_ (80/20; 1 bar) and showed that increasing temperatures (from 5 to 20 °C) may increase the proportion of CH_4_ produced by hydrogenotrophs. Hence highlighting the necessity to better understand the effect of warming on the different methanogenic pathways and communities. Moreover, the results obtained could have been affected by the artificial concentration of acetate used in the experiments as discussed before. Then, further work is necessary to investigate the response to temperature in real conditions. Our results showed that members of *Methanosarcina* and *Methanosaeta* can outcompete other genera at all temperatures, showing an adaptation to both high and low temperatures. As discussed, these genera are widely reported inhabiting cold environments, in this psychrotolerant range (5 – 20 °C). Considering natural temperatures (*in situ*) measured from each ecosystem, our results showed the capacity of methanogens to cope with worse warming predictions; with the concomitant effect of increasing the CH_4_ production and the greenhouse effect associated with CH_4_.

## 5. CONCLUSION

Alaska American state as the Arctic region is considered a susceptible spot to be drastically changed in response to the current global warming phenomenon. The present work consisted in evaluating acetoclastic methanogenic potential from several lake and wetland soil ecosystems (not reported regarding archaeal diversity so far) at increasing temperature, through acetate amended incubations. The results corroborate with current works that increasing temperature has a direct influence and can boost the CH_4_ production in natural environments. In the archaeal communities, different patterns were observed after incubation to higher temperatures, with an increase in *Methanosarcina* abundance for most of the samples (lake sediments and wetland soils) and a persistence of *Methanosaeta* in one of the lakes tested, showing the adaptation of key acetoclastic groups to different temperatures. Besides temperature, organic matter content showed significant correlation with the methanogenic activity.

Archaeal methanogenic diversity reports are scarce in these environments. The present study brings new insights on methanogens from these environments and those involved in acetoclastic methane production in a broader location (center to north) in Alaska, gathering four lakes (Killarney, Otto, Nutella, Goldstream) and two wetland soil (Killarney and Toolik). Our work revealed diverse composition from the studied sites, with strong implications in different biogeochemical processes. Furthermore, the high abundance of Archaea classified as “uncultured” or “not classified” claims for further investigation in order to explore new phyla. The roles and metabolic interaction in metacommunities at increasing temperatures remains not fully elucidated in these underexplored environments, giving a cue to further work on methanogens groups inhabiting these areas. The data gathered here will complement previous works performed in these environments in Alaska (Sepulveda-Jauregui et al., 2015, Cruz et al., 2017) in order to better understand the complex process of CH_4_ cycle in climate change scenario in Alaska.

