## Supplementary Material for "Temperature increase affects acetate-derived methane production in Alaskan lake sediments and wetland soils"

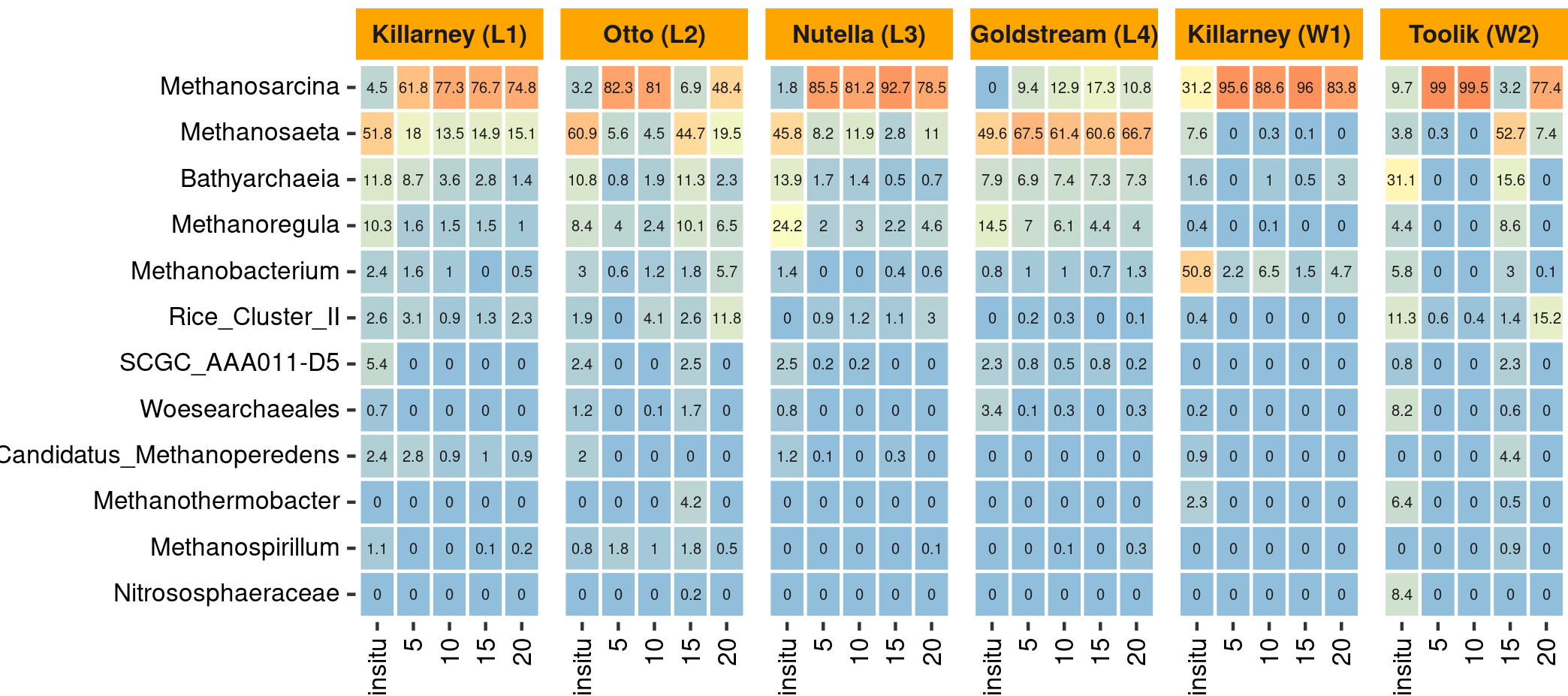

**Figure S1.** Heat map showing the relative abundance of predominant genera according to the archaeal 16S rRNA gene sequence analysis in lake sediment Killarney (L1), Otto (L2), Nutella (L3) and Goldstream (L4) and top layers from wetland soil samples Killarney (W1T) and Toolik (W2T).

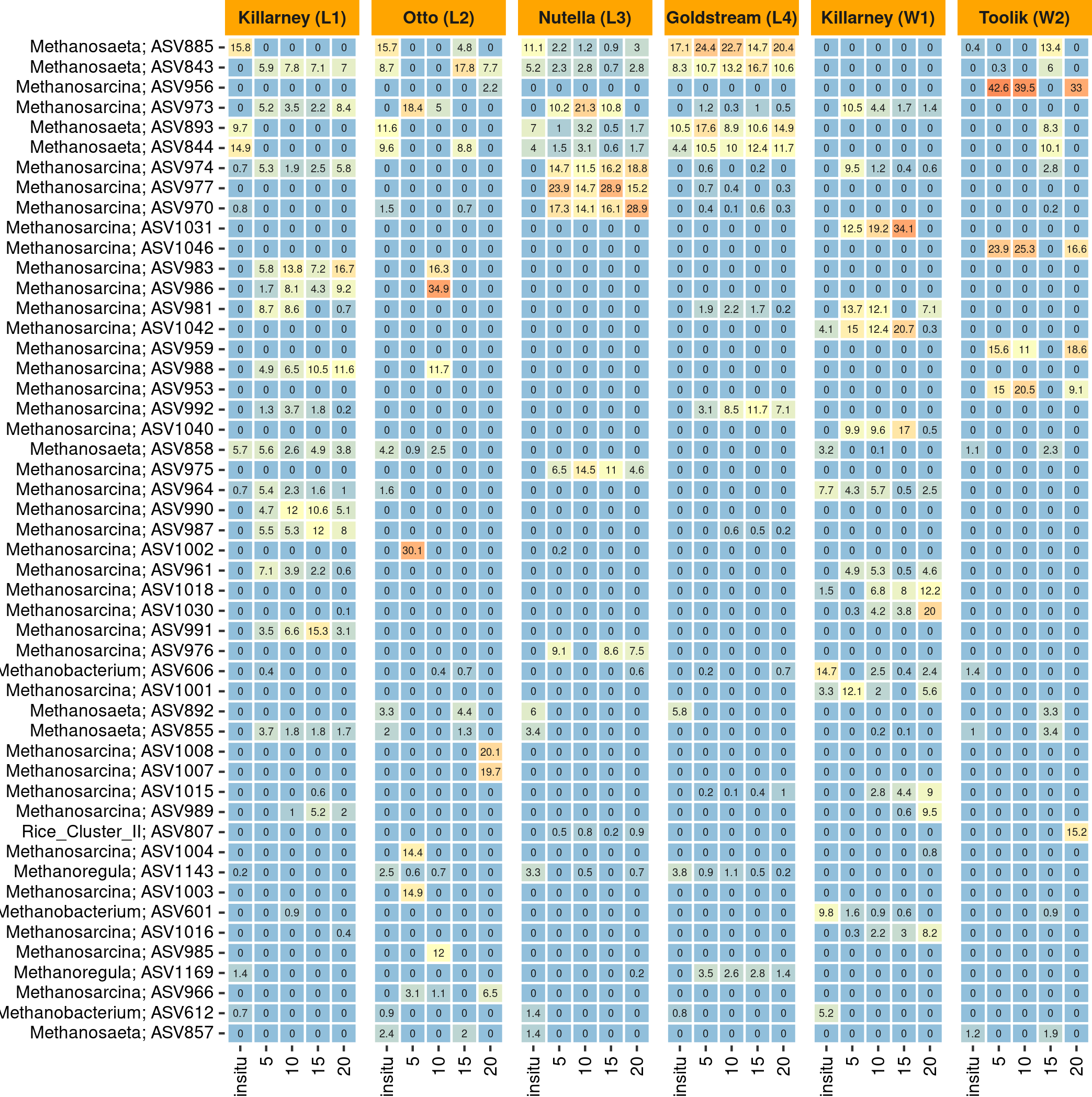

**Figure S2:** Fifty most abundant ASVs classified within the genera *Methanosarcina* and *Methanosaeta* in lake sediment and wetland soil samples, previous and after incubation at four different temperatures.

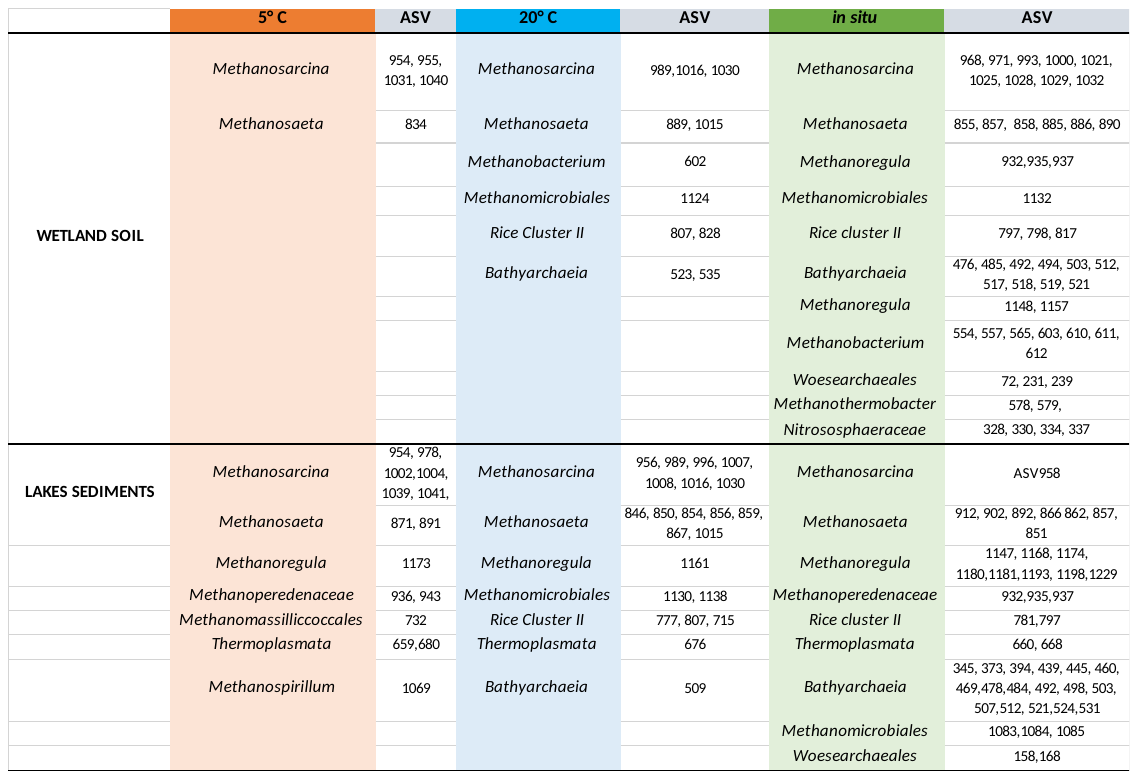

**A**

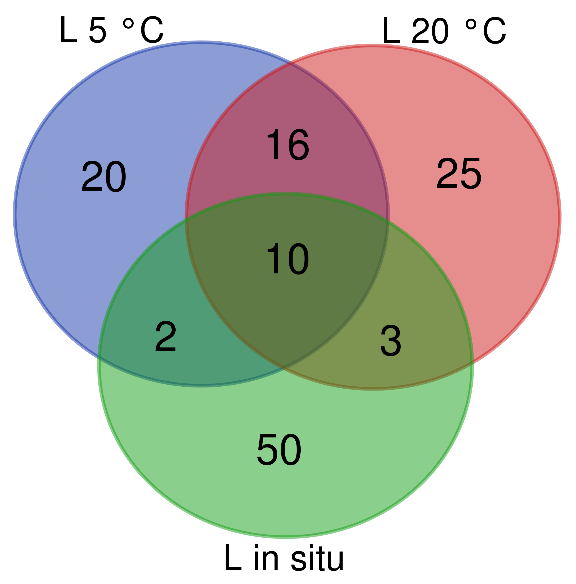

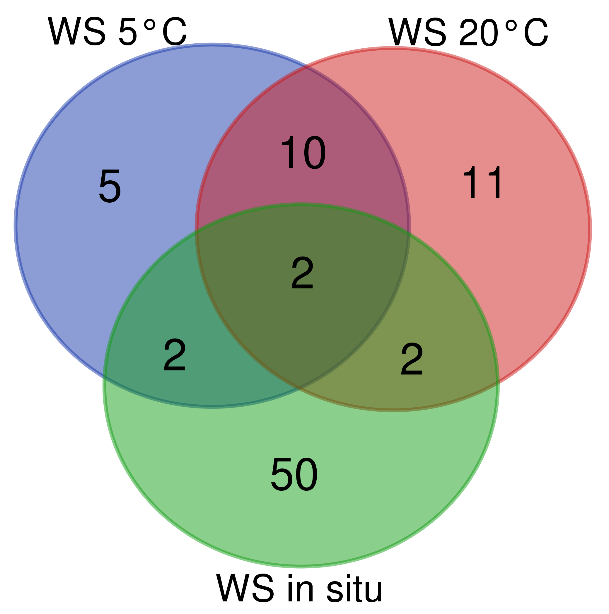

**A**

**B**

**Figure S3**: Venn diagram showing the number of shared ASVs between samples taken in natural condition (in situ), and after incubation at the lowest and highest temperatures (5 and 20°C) in **A)** lake sediments (L) and **B)** wetland soils samples (WS).

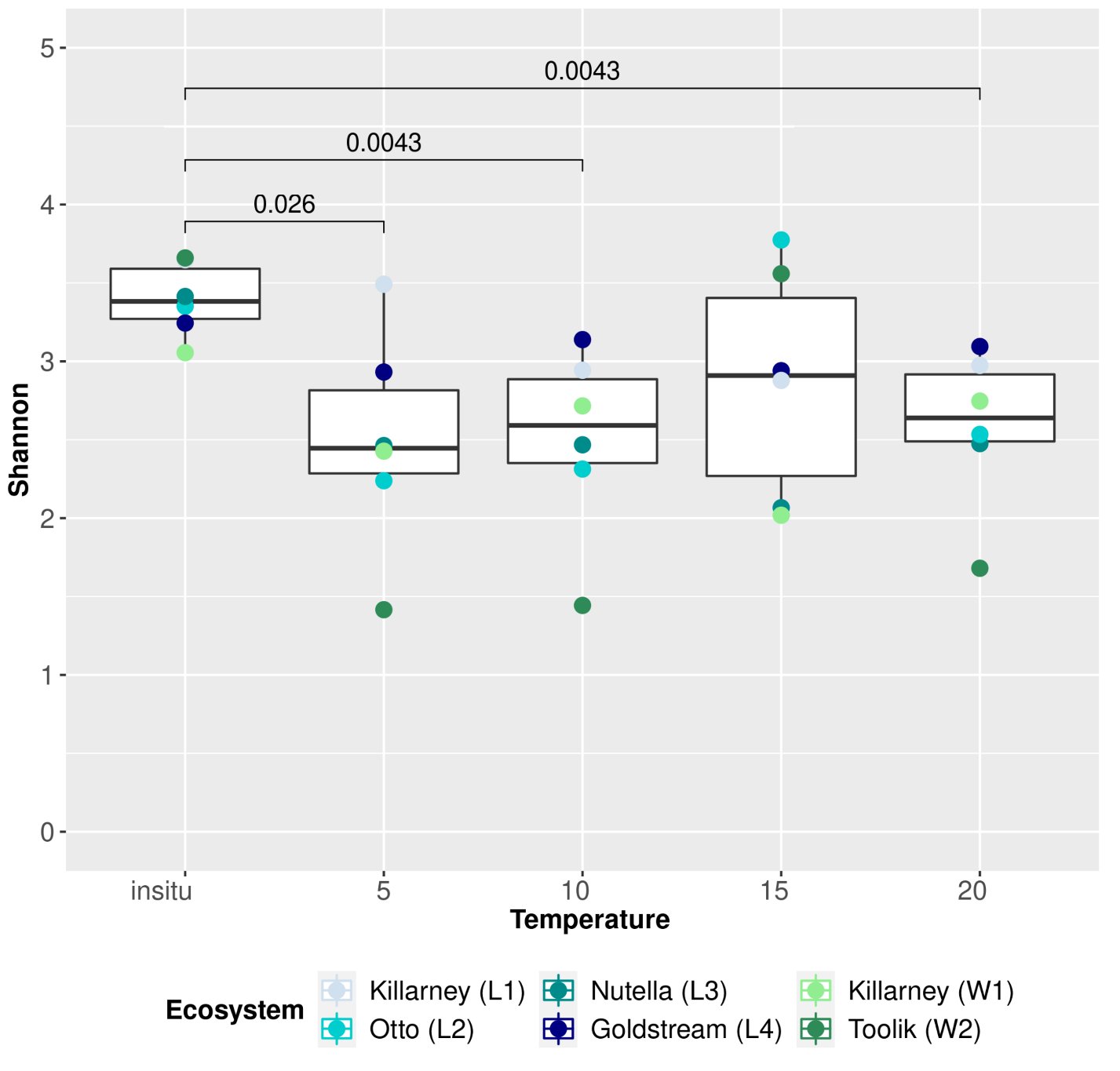

**Figure S4:** Shannon index depicting alpha diversity in lake sediments (Killarney, Otto, Nutella and Goldstream) and wetland soils (Killarney and Toolik) in natural condition (*in situ*) and at four different temperatures (5, 10, 15 and 20 °C).

**Table S1**: qPCR primers

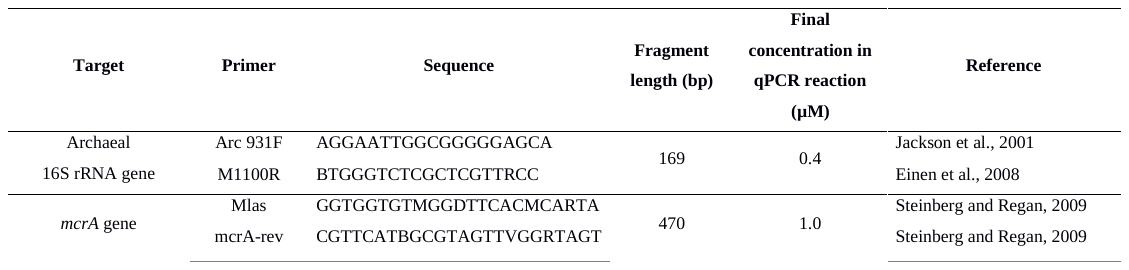

**Table S2**: ASVs found in wetland soils and lake sediments samples in incubations at 5°C and 20°C and environmental (*in situ*) conditions. **A)** ASVs occurring solely at 5 °C, 20 °C and *in situ*; **B)** ASV shared among conditions

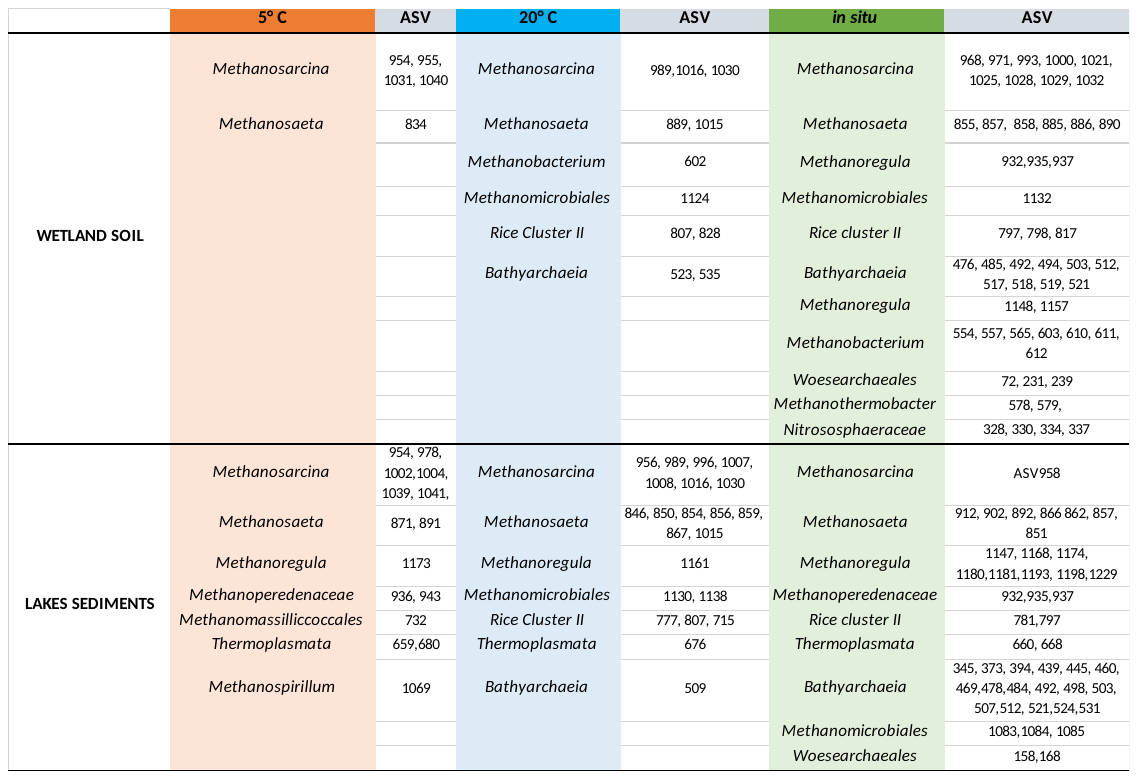

**A**

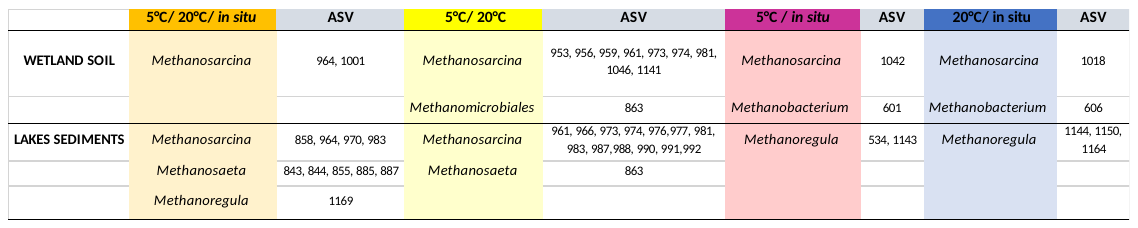

**B**

**Additional Material and Methods**

**Abundance of targeted *mcrA* and *16SrRNA* genes by qPCR**

The pairs of primers used for the ribosomal gene 16S from archaeal group and functional gene *mcrA* are described in **Table 3**. For archaeal 16S rRNA gene, the thermal program used was: 95°C for 3 min, 40 cycles of denaturation 10 s at 95°C, annealing 20 s at 62°C, extension 30 s at 72°C (with a plate read). The cycling was followed by a denaturation at 95°C for 30 sec and a melting curve analysis from 65 to 95°C. Standard curve was run for each experiment, prepared from 10-fold dilutions of archaeal 16S rRNA gene amplified from Clone Arch 21-10 (KT351355) cloned in pGEM®-T vector (Promega). The qPCR efficiencies were 92-100% and correlation coefficients (R²) were always higher than 0.998. This quantification technique allowed to accurately quantify until 4.02 copies of archaeal 16S rRNA gene with a detection limit of 1.33 gene copies.

For *mcrA* gene, the thermal program used was: 95°C for 3 min, 40 cycles of denaturation 10 s at 95°C, annealing 20 s at 55°C, extension 30 s at 72°C and a plate read (8 s at 80°C) to deal with highly degenerated primers. The cycling was followed by an additional extension step 8 min at 72°C, a denaturation at 95°C for 30 sec and a melting curve analysis from 65 to 95°C. Standard curve was run for each experiment, prepared from 10-fold dilutions of *mcrA* gene amplified from *Methanosarcina barkeri CM1* (CP008746) cloned in pGEM®-T vector (Promega). The qPCR efficiencies were 90-99% and correlation coefficients (R²) were always higher than 0.99. This quantification technique allowed to accurately quantifying until 8.48 copies of *mcrA* gene with a detection limit of 2.80 gene copies.

**Abundance of *mcrA* and *16SrRNA* genes by qPCR**

The pairs of primers used for the ribosomal gene 16S from archaeal group and functional gene *mcrA* are described in **Table 3**. For archaeal 16S rRNA gene, the thermal program used was: 95°C for 3 min, 40 cycles of denaturation 10 s at 95°C, annealing 20 s at 62°C, extension 30 s at 72°C (with a plate read). The cycling was followed by a denaturation at 95°C for 30 sec and a melting curve analysis from 65 to 95°C. Standard curve was run for each experiment, prepared from 10-fold dilutions of archaeal 16S rRNA gene amplified from Clone Arch 21-10 (KT351355) cloned in pGEM®-T vector (Promega). The qPCR efficiencies were 92-100% and correlation coefficients (R²) were higher than 0.998. This quantification technique allowed to accurately quantify until 4.02 copies of archaeal 16S rRNA gene µL^-1^ with a detection limit of 1.33 gene copies per µL^-1^.

For *mcrA* gene, the thermal program used was: 95°C for 3 min, 40 cycles of denaturation 10 s at 95°C, annealing 20 s at 55°C, extension 30 s at 72°C and a plate read (8 s at 80°C) to deal with highly degenerated primers. The cycling was followed by an additional extension step 8 min at 72°C, a denaturation at 95°C for 30 sec and a melting curve analysis from 65 to 95°C. Standard curve was run for each experiment, prepared from 10-fold dilutions of *mcrA* gene amplified from *Methanosarcina barkeri CM1* (CP008746) cloned in pGEM®-T vector (Promega). The qPCR efficiencies were 90-99% and correlation coefficients (R²) were higher than 0.99. This quantification technique allowed to accurately quantifying until 8.48 copies of *mcrA* gene µL^-1^ with a detection limit of 2.80 gene copies µL^-1^ .
